# ChAHP2 and ChAHP control diverse retrotransposons by complementary activities

**DOI:** 10.1101/2024.02.05.578923

**Authors:** Josip Ahel, Aparna Pandey, Michaela Schwaiger, Fabio Mohn, Anja Basters, Georg Kempf, Aude Andriollo, Lucas Kaaij, Daniel Hess, Marc Bühler

**Affiliations:** Friedrich Miescher Institute for Biomedical Research, Maulbeerstrasse 66, 4058 Basel, Switzerland; University of Basel, Petersplatz 10, 4003 Basel, Switzerland; Swiss Institute of Bioinformatics, 4058 Basel, Switzerland

## Abstract

Retrotransposon control in mammals is an intricate process that is effectuated by a broad network of chromatin regulatory pathways. We previously discovered ChAHP, a protein complex with repressive activity against SINE retrotransposons, composed of the transcription factor ADNP, chromatin remodeler CHD4, and HP1 proteins. Here we identify ChAHP2, a protein complex homologous to ChAHP, wherein ADNP is replaced by ADNP2. ChAHP2 is predominantly targeted to ERVs and LINEs, via HP1β-mediated binding of H3K9 trimethylated histones. We further demonstrate that ChAHP also binds these elements in a mechanistically equivalent manner to ChAHP2, and distinct from DNA sequence-specific recruitment at SINEs. Genetic ablation of ADNP2 alleviates ERV and LINE1 repression, which is synthetically exacerbated by additional depletion of ADNP. Together, our results reveal that the ChAHP and ChAHP2 complexes function to control both non-autonomous and autonomous retrotransposons by complementary activities, further adding to the complexity of mammalian transposon control.

## INTRODUCTION

Retrotransposons make up a substantial proportion of mammalian genomes and present a potential threat due to their ability to amplify and insert into new genomic loci. The complement of retrotransposons is diverse both in terms of sequence and origin, making their regulation mechanistically challenging^1,2^. To overcome this diversity, cells employ a vast number of different specificity factors that guide a variety of interlinked chromatin-modulating activities. One of the best described examples of this is the repression of endogenous retroviruses (ERVs). Here, the long terminal repeat (LTR) region of the ERV, which normally acts as a promoter, is recognized by sequence-specific transcription factors (TFs) from the Krüppel associated box zinc finger (KRAB-ZFP) protein family^3–5^. These TFs then recruit the co-repressor TRIM28, which in turn interacts with SETDB1, guiding the deposition of H3K9me3 and establishing a transcriptionally silent state^3,6–9^. Additional repressive activity is provided by the HUSH complex and several other mechanisms^10–12^. Firstly, histone deacetylation and the removal of activating histone methylations disfavor transcription^13,14^. Secondly, the underlying DNA is extensively methylated, further repressing the locus^7,9,15,16^. In addition, several chromatin remodelers have been suggested to take part in ERV repression, including ATRX-DAXX, the NuRD complex, MORC3 and SMARCAD1^14,17–19^. Together, these processes are thought to promote chromatin compaction, reinforced by HP1 proteins^20,21^. Similar mechanisms have been proposed to repress non-LTR containing long interspersed elements (LINEs) such as LINE1, albeit with different hierarchies of importance^12,22–24^. Like ERVs, LINE1 retrotransposons are autonomous, and their gene products are required by the non-autonomous short interspersed nuclear elements (SINEs) for retrotransposition^25–27^. This can induce mutations that have been found to cause a variety of heritable and somatic diseases^28–32^.

Retrotransposon families such as SINE retrotransposons, which are derived from tRNAs or 7SL RNAs, can have vastly different characteristics and their regulation is less well understood^25–27,33–35^. We recently discovered part of their repression is conferred by the ChAHP complex, which unites the transcription factor ADNP, chromatin remodeler CHD4, and HP1 proteins into a stably associated module^36,37^. Genetic removal of ADNP allows increased expression of evolutionarily less divergent SINE B2 elements in mouse ES cells (mESCs), accompanied by an increase in chromatin accessibility and CTCF binding^36,37^. Phenotypically, defects in *in vitro* differentiation and mouse development have been observed upon *adnp* knock out, resulting in a penetrant embryonic lethality phenotype by day E9.5^36,38^. In humans, even heterozygous partial truncations of ADNP that abrogate interactions with HP1 proteins result in a severe autism-spectrum syndrome typified by developmental defects, compromised function of multiple organ systems, and intellectual disability^39^. Although their underlying molecular deficiency is unclear, these striking phenotypes highlight the importance of identifying hitherto unknown retrotransposon-associated chromatin regulators.

Here we report the discovery of ChAHP2, a protein complex with similar composition to ChAHP. ChAHP2 is defined by ADNP2, a paralogue of ADNP widely present in vertebrates^40^. ChAHP2 chromatin binding specificity is distinct from ChAHP, predominantly associating with ERV and LINE1 retrotransposons via HP1β-mediated binding of H3K9 trimethylated histones. Through complementary activities, the ChAHP and ChAHP2 complexes control a wide variety of molecularly disparate retrotransposons, including SINEs, LINEs, and ERVs.

## RESULTS

### ADNP2 interacts with HP1β and CHD4 to form a distinct ChAHP2 complex

The domain architecture of ADNP2 resembles that of ADNP, with nine zinc fingers distributed in two N-terminal clusters and one C-terminal homeodomain. These domains are the only regions showing overall high conservation, while the putatively poorly structured linker region between the zinc finger clusters, and the C-terminal unstructured region of ADNP are not well conserved between the two paralogues (Figure 1A). Notably, although the zinc-coordinating residues and overall predicted fold of the zinc fingers are conserved, the residues that normally determine sequence specificity of zinc fingers vary between ADNP and ADNP2 (Figure 1B, Figure S1A). By contrast, the C-terminal HP1 interaction motif (PxVxL)^41,42^ is well conserved. Moreover, ADNP2 is co-purified in CHD4 immune precipitations (IPs)^36^, suggesting that the regions critical for both CHD4 and HP1β/γ interactions are conserved between the two paralogues. These observations prompted us to hypothesize that ADNP2 can form an alternative ChAHP complex, with potentially different properties. To test this, we introduced an affinity tag consisting of Avi-3xFLAG to the C-terminus of ADNP2 in mouse ES cells (Figure S1B). We then used these cells to perform affinity purifications with streptavidin and analyzed the samples by liquid chromatography-mass spectrometry (LC-MS), with the parental cell line serving as a negative control. The bait protein (ADNP2) was strongly enriched in the ADNP2 IPs compared to control, co-purifying CHD4 and HP1β as the top two significantly enriched interactors (Figure 1C). Several other proteins were identified as significantly enriched, but their enrichment was overall lower (Table S1). This group included abundant proteins which commonly co-purify in IPs of DNA-binding factors, such as PARP1, hinting that these may be experimental artefacts^43^. Overall, this experiment shows that ADNP2 interacts with both HP1β and CHD4 in mouse ES cells. Such an interaction pattern is reminiscent of the ChAHP complex, with the notable difference that ADNP2 preferentially binds HP1β over HP1γ (Figure 1D)^36^. Importantly, no ADNP was co-purified with ADNP2 and vice-versa, suggesting that their interaction networks are separate. To further probe the biochemical independence of ADNP and ADNP2, we evaluated whether the composition of either complex is affected by removal of the other. To that end, we knocked out *adnp2* from endogenously edited cells expressing ADNP^FKBP-3xFLAG-Avi^ (Figure S1C) and knocked out *adnp* from cells expressing ADNP2^Avi-3xFLAG^. We screened for successful gene deletion by PCR and confirmed loss of mRNA either by exon-spanning RT-qPCR or RNA sequencing (Figure S1D, Figure S1E). No significant difference in core subunit pulldown efficiencies was apparent between the WT or KO conditions for either ADNP or ADNP2 (Figure S2). This indicates that competition between ADNP and ADNP2 for subunit binding does not have a major effect on complex assembly.

**Figure 1.**
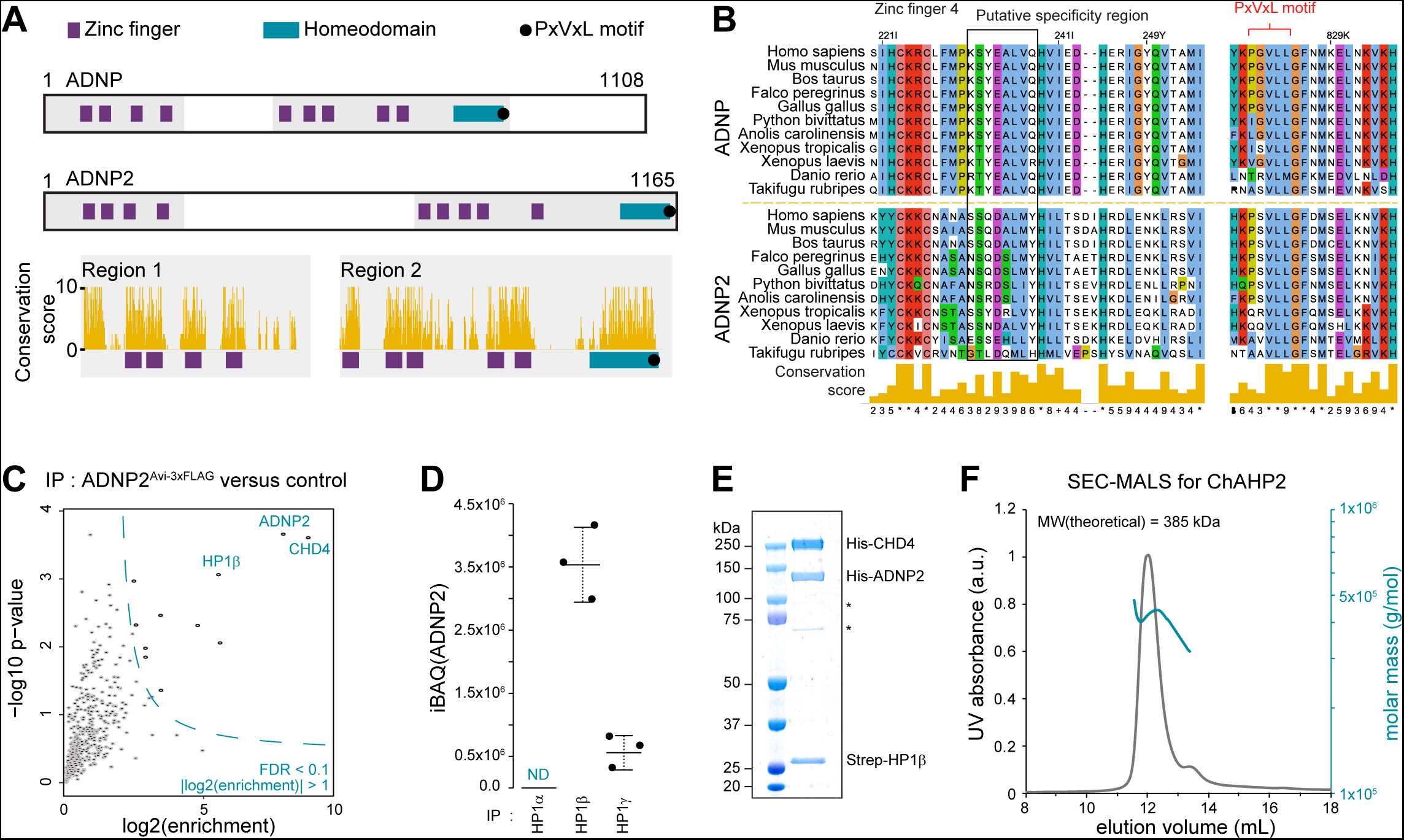
ADNP2 interacts with HP1β and CHD4 to form a ChAHP2 complex. **(A)** Protein architecture schematics of ADNP and ADNP2, drawn to scale. The amino-acid conservation score (Clustal/JalView) is given for the highlighted regions. **(B)** Predicted ADNP or ADNP2 orthologue sequences from species representing different vertebrate classes were aligned using Clustal omega^57^, ordered by species, and visualized with JalView^58^. The conservation scores were scaled and used in Figure 1A. Alignment excerpts for the regions containing Zinc Finger 4 (counting from the N terminus) and PxVxL motif are displayed. The putative region responsible for sequence specificity of the zinc finger is highlighted. Alignment excerpts covering all other Zinc Fingers can be found in Figure S1A. **(C)** Cells expressing endogenously edited ADNP2^Avi-3xFLAG^ or parental untagged cells were subjected to immune precipitation with Streptavidin and analyzed by proteomics (n=3). Dashed line represents the significance cutoff at FDR = 0.1 and log2Enrichment > 1. **(D)** Abundance of proteins from the HP1 proteins recovered in the experiment described in (C), expressed as iBAQ values. **(E-F)** His-ADNP2 and Strep-HP1β were co-expressed, and His-CHD4 expressed separately using baculovirus transductions in insect cells before lysing the mixed cell pools and pulldown against Strep, followed by electrophoresis and Coomassie staining **(E)** and SEC-MALS **(F)**.

Finally, to directly confirm that ADNP2, HP1β and CHD4 form a stable complex, we performed *in vitro* reconstitutions. First, we co-expressed His-ADNP2 and Strep-HP1β in insect cells and expressed His-CHD4 in separately transduced cells, before mixing the two cell pools for lysis and Streptactin pulldown. We analyzed these pulldowns using size exclusion chromatography with multi-angle light scattering (SEC-MALS) and Coomassie staining. All three proteins were detected in one elution peak, with an estimated molecular weight consistent with an ADNP2-CHD4-HP1β complex (Figure 1E, Figure 1F). Together, these data demonstrate that ADNP2, CHD4 and HP1β form an independent *bona fide* complex, which we refer to as ChAHP2.

### ChAHP2 binds retrotransposons

We next performed ChIP-sequencing to characterize the chromatin localization of ADNP2. Peak calling revealed 6315 regions with significant ADNP2 enrichment over input chromatin (Table S2). These regions also showed enriched binding of CHD4 and HP1β in previously published datasets, suggesting that ChAHP2 complexes occupy these sites (Figure 2A). Intriguingly, the majority of ADNP2 peaks were found in repeat regions, while few overlapped transcription start sites (TSS) (Figure 2B). Retrotransposons belonging to LTR-containing families, including several classes of ERV elements, were particularly overrepresented when compared to a randomized peakset of equal properties (Figure 2C, Figure 2D). In addition, one subfamily of LINE1 elements was overrepresented (Figure 2D). We further supplemented these analyses with a peak-agnostic, repeat-family-based approach. This agreed well with the initial analysis, showing enrichment over input at the repeat classes identified by peak calling, with both the internal sequence of ERVs and their associated LTR sequences exhibited enrichment in ADNP2 ChIP signal (Figure 2E, Figure S3A). Finally, we additionally determined ADNP2 ChIP signal distribution along repeat elements using mapping to repeat consensus sequences, which showed similar results to the other analyses (Figure 2F, Figure S3B). Together, these data indicate that ADNP2 binds several classes of retrotransposons both internally and at their terminal sequences.

**Figure 2.**
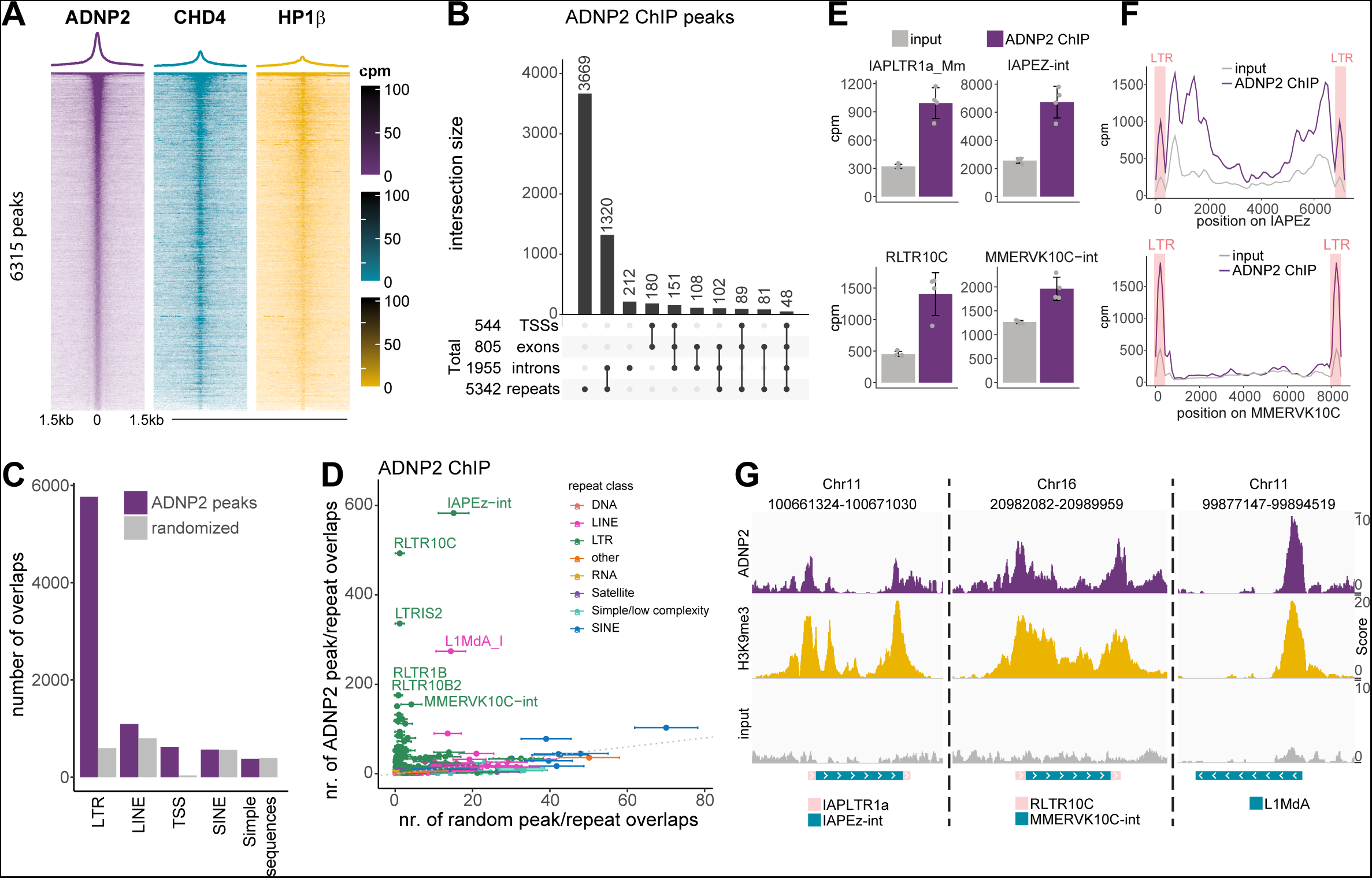
ChAHP2 predominantly binds to ERVs and LINEs. **(A)** Heatmaps of ChIP-seq counts normalized to library size centered on summits of ADNP2 peaks (mean, n=4). **(B)** Upset plot of overlaps between ADNP2 peaks and select genomic features (TSSs, exons, introns, repeats). **(C)** Overlaps numbers between repeat annotations and ADNP2 peaks or randomized peakset of equal properties (bootstrapped 100 times, mean ± SD). Simple sequences = simple repeats and low complexity regions. **(D)** Same as (C), but further split by repeat family. **(E)** Summed reads over repeat annotations normalized to library size for input and ADNP2 ChIP-seq (mean ± SD, n=4). **(F)** Consensus mapping traces over repeat annotations normalized to library size for input and ADNP2 ChIP-seq (mean ± SD, n=4). For IAPEz, the IAPEz-int consensus sequence was stitched with IAPLTR1a_Mm at either end, while for MMERVK10C, MMERVK10C-int was stitched with RLTR10C. The position of the stitched LTRs is highlighted. **(G)** IGV genome browser shots of select regions bound by ADNP2.

### ChAHP2 is recruited to chromatin via HP1 binding to H3K9me3

Contrary to the expected behavior of a TF, no DNA sequence motifs accounted for more than 15% of ADNP2-bound sites (Figure S3C). In addition, these motifs were not centrally enriched within ADNP2 peaks, hinting they are not a direct specificity determinant (Figure S3C). Notably, a common feature of ADNP2-bound retrotransposons is the presence of H3K9 trimethylation (Figure 2G)^6,22,44^. Since HP1 proteins are known to bind this histone modification^45,46^, we hypothesized that ChAHP2 is recruited to its targets by HP1β-mediated binding to H3K9me3 histone tails. Indeed, ADNP2 chromatin binding correlates well with H3K9me3 levels genome-wide (Figure 3A). To directly test if HP1 is required for ChAHP2 binding to chromatin, we specifically abrogated the ADNP2-HP1β interaction by introducing point mutations in the PxVxL motif of ADNP2. We verified successful editing and homozygosity by Sanger sequencing (Figure S4A). As expected, wild type ADNP2 was able to pull down HP1β and CHD4, whereas ADNP2^PxVxL^ mutants were unable to bind HP1β above background levels (Figure S4B). Importantly, CHD4 binding was unaffected, confirming the mutations specifically disrupt the ADNP2-HP1β interaction (Figure S4B). Next, we performed ChIP-sequencing for ADNP2 and ADNP2^PxVxL^. Mutations in the PxVxL motif resulted in near complete loss of ChAHP2 binding to H3K9me3 modified target regions while H3K9me3 levels remained unchanged (Figure 3C). These regions almost exclusively overlap LTR or LINE1 elements. Statistical analysis of significantly changing peaks corroborated this initial impression (Figure S4C, Figure S4D). In turn, the peaks with increased binding were promoter-associated, devoid of H3K9me3 and exhibited higher chromatin accessibility (Figure S4E, Figure S4F). This behavior would be consistent with a redistribution of ChAHP2 from heterochromatin to euchromatin when HP1 is removed from the complex. Finally, regions without a significant change exhibited an intermediate accessibility and H3K9me3 profile (Figure S4F). Despite not being identified as statistically significant, ADNP2 binding was still overall reduced in this group (Figure S4G). These observations provide evidence that chromatin binding at the majority of ChAHP2 targets is dependent on HP1.

**Figure 3.**
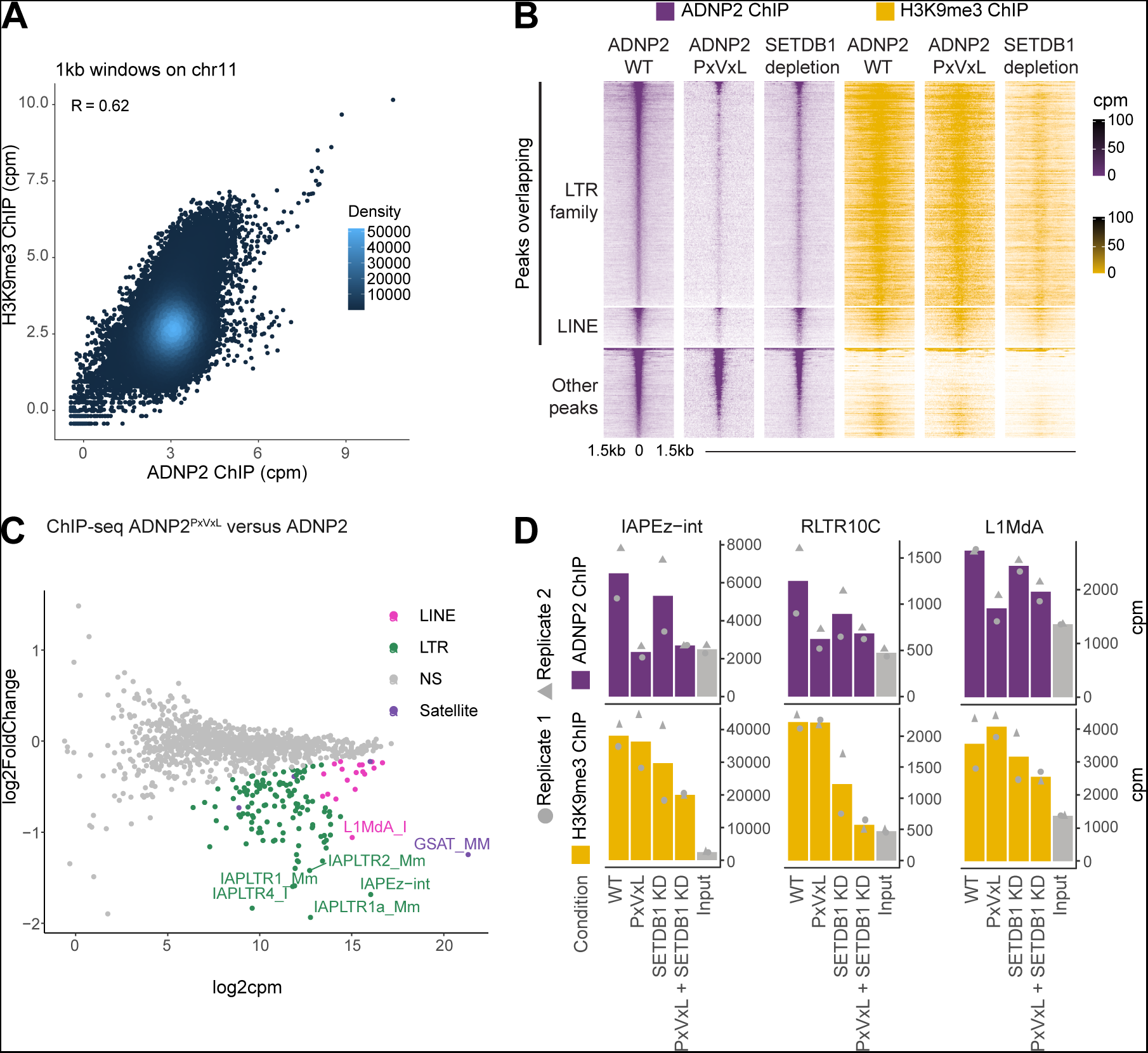
ChAHP2 binds heterochromatin in an HP1- and H3K9me3-dependent manner. **(A)** Comparison between ADNP2 and H3K9me3 ChIP-seq signal in 1kb genomic bins across chromosome 11. **(B)** Cells expressing WT or PxVxL mutated ADNP2^Avi-3xFLAG^, before or after SETDB1 depletion were analyzed by ChIP sequencing. Average ChIP cpm displayed (n=2). **(C)** Differential binding analysis of summed reads over repeat annotations normalized to human spike-ins comparing ADNP2^PxVxL^ ChIP and WT controls (n=2). **(D)** Summed reads over repeat annotations normalized to human spike-ins for input and a series of ChIP samples as indicated (mean, n=2, replicates annotated). ADNP2 and H3K9me3 ChIPs were done simultaneously and from the same material.

We next sought to test whether ChAHP2 binding depends on the presence of H3K9me3. To do this, we introduced a 2xHA-FKBP12^F36V^ degron tag at the N-terminus of endogenous SETDB1 – the methyltransferase responsible for depositing H3K9me3 at LTR elements (Figure S5A)^6,47^, and confirmed the efficacy of SETDB1 depletion by western blotting (Figure S5B). We then generated ADNP2 and H3K9me3 ChIP-seq data in untreated and SETDB1-depleted conditions. Consistent with its described roles, SETDB1 depletion resulted in a reduction of H3K9me3 at LTR elements, and a modest reduction at minor target regions such as LINE1 elements (Figure 3B, Figure 3D)^6,7,24^. Tracking these changes, ADNP2 binding was reduced at LTRs, and to a lesser extent at LINE elements (Figure 3B). This reduction was smaller than the effect of the PxVxL mutation, likely due to incomplete loss of H3K9me3 at these elements. Supporting this interpretation, the change in ADNP2 binding strongly correlated with the change in H3K9me3 levels across the entire dataset (Figure S5C).

To further quantify the relationship between ChAHP2 and H3K9me3, we analyzed the HP1 decoupling and SETDB1 depletion data using repeat-family-wide and consensus mapping approaches. First, we performed a differential binding analysis between ADNP2 and ADNP2^PxVxL^ mutant ChIPs across all transposon families (Figure 3C). This revealed a significant binding reduction in the PxVxL mutant for many LTRs and LINE1s. Similarly, binding to satellite repeats (GSAT_MM) was reduced, suggesting binding to non-transposon heterochromatin is also HP1-dependent. Consistent with the peak-based analyses (Figure S4C, Figure S4D, Figure S4E), no transposon class exhibited significantly increased binding (Figure 3C). A focused look at representative LTR elements and L1MdA confirmed that ADNP2 binding dropped to input levels at H3K9 trimethylated transposons in the HP1−deficient mutant (Figure 3D, Figure S5D). A reduction was also apparent upon SETDB1 depletion, again smaller in scale. Finally, there was no additional reduction in ADNP2 ChIP signal when SETDB1 was depleted in the ADNP2^PxVxL^ background (Figure 3D), suggesting the two perturbations exert their effect via a shared molecular axis. Overall, these data demonstrate that HP1β-mediated binding of H3K9me3 nucleosomes targets ChAHP2 to heterochromatin.

### ChAHP partially co-localizes with ChAHP2 at retrotransposons

These chromatin binding characteristics appear overall distinct from ChAHP, which has been shown to predominantly bind SINE elements^36,37^. We see little evidence of ADNP2 binding at ADNP-associated SINE elements, with few ADNP2 peaks overlapping these transposons, and very little ADNP2 signal at ADNP peaks (Figure 2C, Figure 2D, Figure 4A, Figure 4B, Figure 4E). Conversely, some ADNP does colocalize with ADNP2, specifically at regions marked by H3K9me3, though this binding is less strong than on SINEs (Figure 4A, Figure 4B). Indeed, we observe ADNP peaks (Table S3) at select heterochromatic repeat regions more frequently than a randomized peak set (Figure S6A). ADNP has been previously reported to bind to H3K9me3 modified chromatin and pericentromeric heterochromatin in an HP1-dependent manner^36,42^, prompting us to assess whether ADNP is targeted to ADNP2-bound sites via the same mechanism. To do this, we generated ADNP^PxVxL^ motif point mutants (Figure S6B) and performed ChIP sequencing. Abrogating the HP1 interaction in this way resulted in loss of ADNP signal specifically at H3K9 trimethylated transposons, including LTR and LINE1 families which are bound by ADNP2 (Figure 4C, Figure 4E). Binding to satellite repeats was also significantly reduced, orthogonally validating previously published imaging data (Figure 4C)^42^. A focused look at representative heterochromatinized repeats confirmed that ADNP binding at these sites drops to near background levels after HP1 decoupling (Figure 4D, Figure S6C). Conversely, binding to SINE elements is increased under these conditions (Figure 4B, Figure 4C, Figure 4D, Figure S6C). This striking observation indicates that ChAHP features two chromatin binding modes: one via HP1 and H3K9me3 to heterochromatin analogous to ChAHP2, and a second H3K9me3-independent mechanism through sequence specific recruitment to euchromatic SINEs^36,42^. Thus, these analyses demonstrate that ChAHP and ChAHP2 have different combinations of chromatin binding specificities, with a notable overlap at heterochromatin.

**Figure 4.**
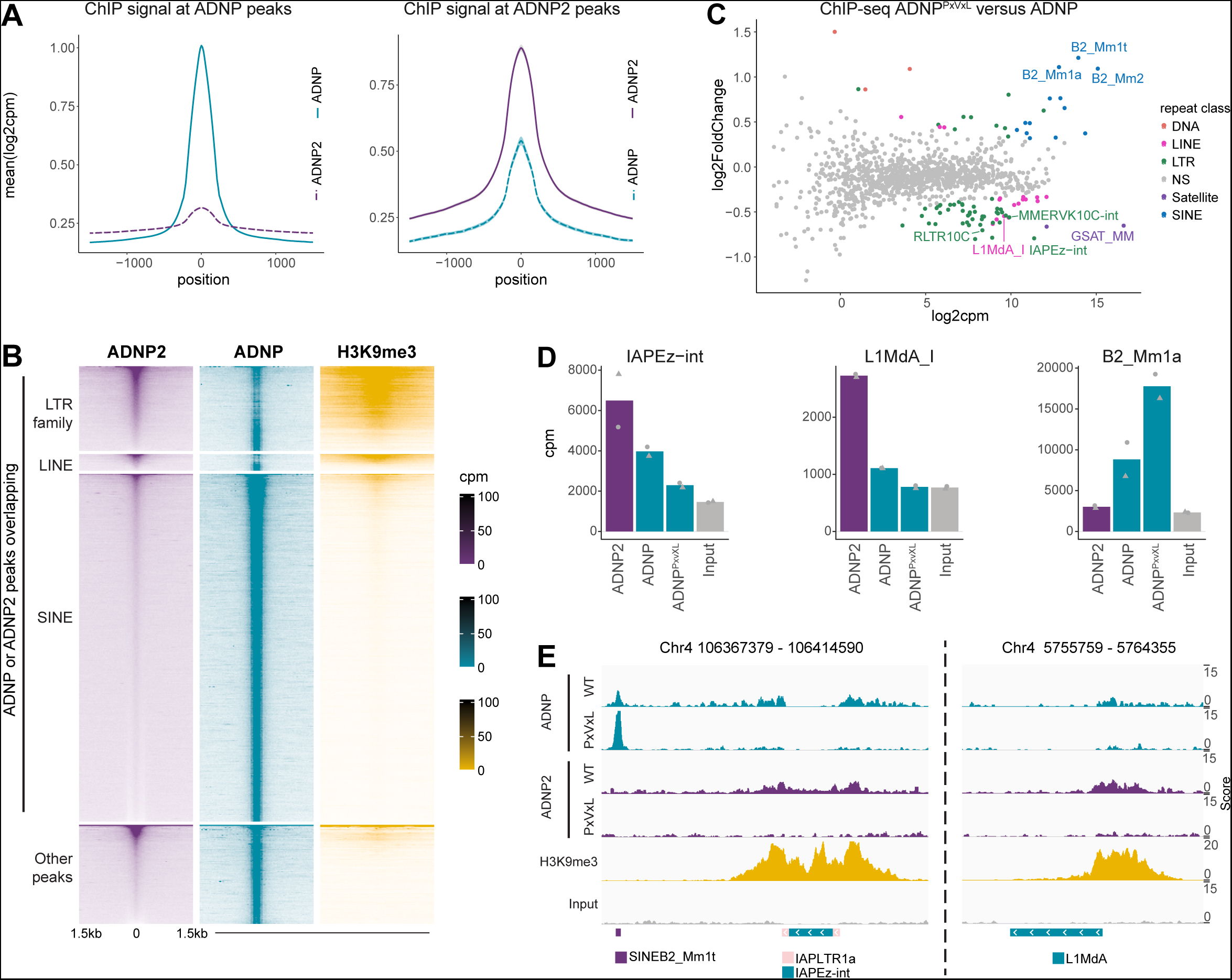
ChAHP and ChAHP2 colocalize at heterochromatin. **(A)** Metaplots of ChIP-seq counts normalized to library size over ADNP peaks or ADNP2 peaks as indicated (mean ± SD, n=2). **(B)** Waterfall plot for ChIPs, centered on a concatenated set of ADNP and ADNP2 peak summits, and split by overlap with select repeat annotations as indicated (mean, n=2). **(C)** Differential binding analysis of summed reads over repeat annotations normalized to library size comparing ADNP^PxVxL^ ChIP and WT controls (n=2). **(D)** Summed reads over repeat annotations normalized to library size for input and a series of ChIP samples as indicated (mean, n=2, replicates represented as points). **(E)** IGV genome browser shots for select regions for ChIPs and input as indicated.

### Repressive activities of ChAHP and ChAHP2 partially overlap at LINE1 and LTR retrotransposons

The chromatin binding characteristics of the two ChAHP complexes prompted us to assess their individual and combined contribution to transposon silencing. We attempted to generate *adnp/adnp2* double KO cells, but were unable to obtain homozygous clones. To circumvent this issue, we generated cells that allow inducible degradation of ADNP in an *adnp2^−/−^* background. Approximating a constitutive KO situation, we induced ADNP degradation in *adnp2^−/−^*cells continuously over 14 days of passaging and confirmed the efficacy of degradation by western blotting (Figure S7A). At the RNA level, consistent with previous data, several dozen genes were either up- or downregulated upon removal of ADNP (Figure S7B, Table S4)^36^. Genetic KO of *adnp2* caused misregulation of several hundred genes, with a slight tendency for upregulation (Figure S7B, Table S4). Depletion of ADNP in *adnp2^−/−^* cells resulted in misregulation of several hundred genes in addition to those already observed when only ADNP2 was removed, revealing a synthetic effect (Figure S7B, Table S4). None of the changing gene categories showed strong and significant patterns either in terms of GO term enrichment (FigureS7C), distance between the promoter and the nearest ADNP/ADNP2 peaks (Figure S7D), or direct association between the peaks and promoters (Hypergeometric test, p < 0.01). Therefore, ChAHP and ChAHP2 likely do not regulate these genes directly.

We next explored whether any repetitive elements were differentially expressed under the same conditions (Figure 5A, Table S5). Consistent with previous findings, only SINE B2 elements were significantly upregulated upon ADNP depletion (Figure 5B)^37^. Conversely, nine repeat families were significantly upregulated in *adnp2^−/−^*cells (FDR < 0.05, |Log2FoldChange| > 0.9), while two were downregulated (Figure 5A). All these differentially expressed repeats belonged to the LTR class of retrotransposons, including families identified as ADNP2-bound in ChIP-sequencing, such as those corresponding to components of IAP elements and MMERVK10C. Notably, both the internal sequences of the elements and their associated LTR sequences were upregulated, suggesting that all components of these elements increase in expression. Strikingly, combined removal of ADNP and ADNP2 resulted in magnified upregulation of LTR elements when compared to ADNP or ADNP2 removal in isolation. In addition, two LINE1 element subclasses exhibited significant upregulation, with the more prominent being L1MdA. Closer inspection revealed that L1MdA elements are already slightly upregulated upon ADNP2 KO, but the upregulation only reaches significance upon removal of both ADNP and ADNP2 (Figure 5D). Synergistic regulation was also observed for IAP elements. In contrast, MMERVK10C derepression was exclusively ADNP2-sensitive, despite also being weakly bound by ADNP (Figure 5C). We obtained nearly identical findings when intron-associated repeat elements were excluded from the analyses (data not shown), indicating that the observed effects are not covariates of changes in gene expression. Collectively, these data suggest ChAHP and ChAHP2 contribute to retrotransposon repression with distinct, but partially overlapping specificities.

**Figure 5.**
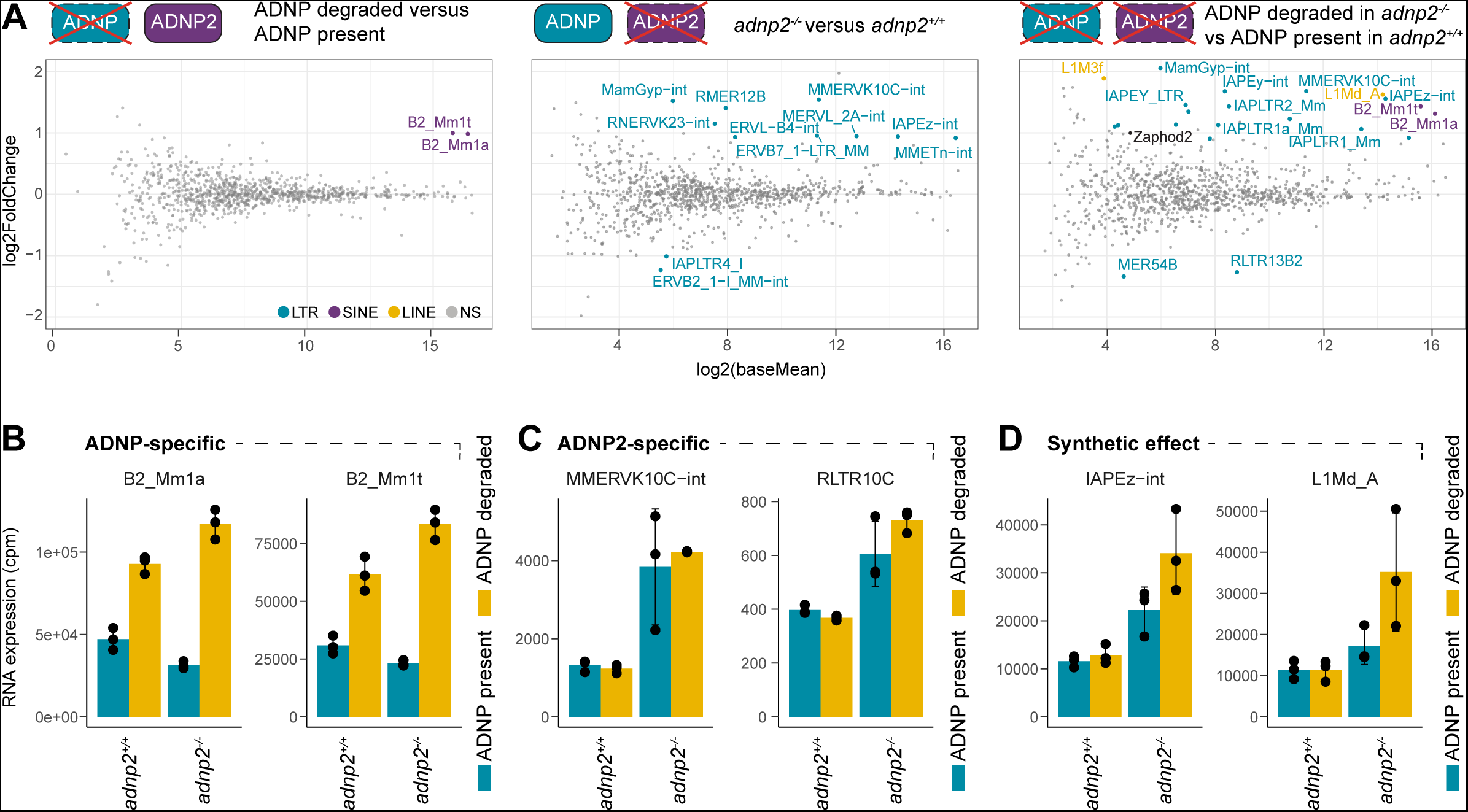
ChAHP and ChAHP2 repress distinct and shared repeat classes. **(A)** Differential expression analysis for repeat families between ADNP/ADNP2 perturbation and the corresponding unperturbed controls (n=3). Significant hits are highlighted with colors (FDR < 0.05, |log2FoldChange| > 0.9). ADNP depletion was performed using 250nM dTAG13 for 14 days, whereas ADNP2 was genetically removed using CRISPR-Cas9. **(B-D)** RNA expression normalized to library size for example repeat classes (mean ± SD, n=3), from the same dataset as in (A), representing classes which are specifically ADNP-dependent **(A)**, specifically ADNP2-dependent **(C)** or cooperatively regulated **(D)**.

## DISCUSSION

Here we have shown that ADNP2, CHD4 and HP1β coalesce to form ChAHP2 – a novel stable protein complex unifying different chromatin regulatory activities. The chromatin binding properties of ChAHP2 are largely distinct from ChAHP in that its predominant recruitment mode is via HP1 and H3K9me3. In contrast to ChAHP, ChAHP2 does not bind euchromatic SINE elements, but is instead targeted to H3K9me3-modified LTR and LINE1 elements. In line with this sequence-agnostic recruitment mechanism, we were unable to define a sequence motif that would broadly explain the distribution and specificity of ADNP2 ChIP-seq signal. It is possible that multiple different sequences can be bound by distinct DNA binding domains within ADNP2 (Zinc fingers or homeodomain) and contribute weak but important specificity towards certain repetitive elements. Thorough *in vitro* biochemistry and potentially structural analyses will be required to address this question in the future. Nevertheless, the observed chromatin binding appears consequential, as removal of ADNP2 results in specific upregulation of a subset of ChAHP2-bound retrotransposons.

We also find a supporting role of ChAHP in regulating these elements. Previous studies had reported weak and promiscuous binding of ChAHP to H3K9me3-modified heterochromatin without a clear functional implication^36,42^. Since ChAHP binding at non-H3K9me3 modified cognate targets is increased upon disrupting HP1 interaction and thus H3K9me3 binding, a substantial fraction of ChAHP is likely associated with heterochromatin. This observation indirectly supports a role for ChAHP in regulation of heterochromatic loci and clarifies the biological requirement for this chromatin binding modality within the complex. This was previously unappreciated because derepression of heterochromatic targets only becomes apparent when both ADNP and ADNP2 are removed. Notably, the magnitude of this derepression is comparable to the contribution of other factors with proposed roles in LTR regulation such as MORC3, ATRX-DAXX, SMARCAD1, m6A methylation, or the recently identified TNRC18^17–19,48–50^. Thus, we surmise that ChAHP and ChAHP2 constitute one part of a multi-component retrotransposon repression machinery.

The most likely mechanism which would confer repressive activities to ChAHP complexes is chromatin remodeling by CHD4. Previously, CHD4 has been shown to contribute to transcriptional silencing via chromatin remodeling activity promoting increased nucleosome densities in a non-positioned manner at its target loci^51–53^. Such a mechanism would conform to the general concept of transposon control, wherein a diverse group of specificity factors guide the activity of less specific chromatin-modifying effectors. This model is typified by the canonical LTR repression system harnessing sequence specific KRAB-ZFPs to guide the activity of H3K9 methyltransferases. There are two notable differences between this system and ADNP/ADNP2. Firstly, chromatin remodeling activity is stably incorporated into the ChAHP complexes, while the KRAB-ZFPs transiently interact with chromatin modifiers and don’t appear to form stable multi-subunit complexes^54^. Secondly, KRAB-ZFPs rapidly evolve and massively multiplied to target newly invading TEs^4,55^, whereas ADNP and ADNP2 are remarkably well conserved since their inception in agnathans. Instead, the recognition of potentially harmful genetic elements in a largely DNA sequence agnostic manner, which is exerted by the H3K9me3 affinity of HP1 proteins within the ChAHP complexes, is conceptually reminiscent of the HUSH complex^10,56^. HUSH recognizes nascent intron-less transcripts – a feature typical for TEs – and chromatin modifications, rather than DNA sequence itself. However, like KRAB-ZFPs, HUSH interacts with chromatin modifiers only transiently. Thus, ChAHP and ChAHP2 possess a unique combination of chromatin binding modes and functional properties, representing previously unknown components of the LTR, LINE1 and SINE repression machineries.

## ACKNOWLEDGEMENTS

We thank the members of the Bühler lab for their constant support and discussions. Special thanks go to Yukiko Shimada and Nathalie Laschet for technical support. We are also grateful to the FMI Functional Genomics facility for library construction and next-generation sequencing. This work was supported by the Novartis Research Foundation, the Boehringer Ingelheim Fonds, and the Swiss National Science Foundation (SNSF, grant 310030_188835).

## AUTHOR CONTRIBUTIONS

J.A. and A.P. generated cell lines, designed and performed most experiments, and analyzed data. M.S. performed the bulk of bioinformatics analyses. J.A. and M.S. generated the figures. F.M. performed and advised experiments, and generated cell lines. A.B. performed protein purifications and *in vitro* biochemistry analyses. G.K. performed SEC-MALS. D.H. acquired and analyzed the mass spectrometry data. A.A. and L.K. generated cell lines. M.B. conceived and supervised the study, and secured funding. J.A., F.M. and M.B. wrote the manuscript. All authors discussed the results and commented on the manuscript.

## FIGURE LEGENDS

**Figure S1, related to Figure 1. (A)** Predicted ADNP or ADNP2 orthologue sequences from species representing different vertebrate classes were aligned using Clustal omega^57^ and visualized with JalView^58^. The conservation scores were scaled and used in Figure 1A. Alignment excerpts for the regions containing Zinc Finger 4 (counting from the N terminus) and PxVxL motif are displayed. The putative region responsible for sequence specificity of the zinc finger is highlighted. **(B)** Sanger sequencing traces of PCR amplicons around the C terminal region of ADNP2 after insertion of the Avi-3xFLAG tag, focused on the site of editing. These cell lines were used in Figure 1B and Figure 1D. **(C)** Sanger sequencing traces of PCR amplicons around the C terminal region of ADNP after insertion of the FKBP-3xFLAG-Avi tag, focused on the site of editing and the STOP codon. These cell lines were used in Figure 1B, Figure 1D and Figure 5. **(D)** RT-qPCR analysis of the indicated RNAs in parental control (cMB581) and derived *adnp^−/−^* clones. Note that ADNP mRNA is undetectable in the knockout clones under these conditions, and SINE B2 RNA is more abundant. These cell lines were used in Figure 1D **(E)** Normalized RNA sequencing counts for *adnp2^+/+^* and *adnp2^−/−^*cells in the ADNP^FKBP-3xFLAG-Avi^ background, used in Figure 1D and Figure 5.

**Figure S2, related to Figure 1. (A)** ADNP2 or ADNP were deleted using genome editing in endogenously edited lines expressing ADNP^FKBP-3xFLAG-Avi^ or ADNP2^Avi-3xFLAG^, respectively, and subjected to immune precipitation against FLAG, followed by proteomics analyses. Comparison of indicated target IP samples as indicated to the untagged controls (n=3). **(B)** Comparison of ADNP IPs between *adnp2^+/+^*and *adnp2^−/−^* or ADNP2 IPs between *adnp^+/+^* and *adnp^−/−^*. C) Intensity values for ChAHP complex components normalised to the bait protein.

**Figure S3, related to Figure 2. (A)** Summed reads over repeat annotations normalized to library size for input and ADNP2 ChIP-seq (mean ± SD, n=4). **(B)** Consensus mapping traces over repeat annotations normalized to library size for input and ADNP2 ChIP-seq (mean ± SD, n=4). The internal consensus sequence was stitched with the 5’ end and 3’ end consensus sequences to generate the plot. The position of the stitched sequence is highlighted. **(C)** Enriched motifs in ADNP2 ChIP peaks, based on MEME-ChIP. For the top 3 motifs by significance, the motif sequence, position, and representation as a percentage of peaks is shown.

**Figure S4, related to Figure 3. (A)** Example Sanger sequencing trace of a PSVLL-PSELT (PxVxL) mutant cell line in the ADNP2^Avi-3xFLAG^ background, used in Figure 3. **(B)** Cells expressing endogenously edited ADNP2^Avi-3xFLAG^ either WT or with a PSVLL-PSELT (PxVxL) mutation were subjected to immune precipitation against FLAG and analyzed by western blotting, with the untagged cell line serving as a control. Inputs and IP excerpts come from the same image, with intervening lanes spliced out. Asterisk denotes the antibody heavy chain band. **(C)** Cells expressing WT or PxVxL mutated ADNP2^Avi-3xFLAG^ were analyzed by ChIP sequencing and normalized to human spike-ins. Differentially bound regions identified using edgeR (n=2). **(D)** Overlap between repeat annotations and peaks with significantly decreased ADNP2 binding upon introduction of the PxVxL mutation, or a randomized region set with equal properties (mean ± SD, boostrapped 100 times, NS = not significant). **(E)** Same as (D), but for regions with significantly increased binding. **(F)** Metaplots of H3K9me3-ChIP and ATAC-sequencing reads of the indicated samples centered on ADNP2 peak summits split into categories based on ADNP2 binding behavior of the PxVxL mutant compared to control. (mean ± SD, n = 2 for H3K9me3, n = 3 for ATAC). **(G)** Same as in (F), but for ADNP2 WT and PxVxL mutant ChIP.

**Figure S5, related to Figure 3. (A)** Example Sanger sequencing traces of the beginning and junction regions in ^2HA-FKBP^SETDB1 cell lines, with an ADNP2^Avi-3xFLAG^ background (parental = cMB580, Figure S1), used in Figure 3. **(B)** Cells were treated with 500nM dTAG13 for 48h before western blot analysis. Asterisk denotes a non-specific band which serves as a loading control. Blots originate from the same image, with intervening lanes spliced out. **(C)** Correlation of changes in H3K9me3 and ADNP2 ChIP-seq signal over ADNP2 peaks upon depletion of SETDB1. **(D)** Consensus mapping traces over repeat annotations normalized to library size for input and ADNP2 or H3K9me3 ChIP-seq split by replicate. The H3K9me3 attrition is more modest in replicate 2, and reflected in a weaker effect on ADNP2 binding. Sequences were stitched as described in Figure 2E.

**Figure S6, related to Figure 4. (A)** Overlaps numbers between repeat annotations and ADNP peaks or randomized peakset of equal properties (boostrapped 100 times, mean ± SD). **(B)** Sanger sequencing traces of PGVLL-PGELT (PxVxL) mutant cell line in the ADNP^3xFLAG-V5^ background, used in Figure 4. **(C)** Summed reads over repeat annotations normalized to library size for input and ChIP samples as indicated (mean, n=2, replicates annotated).

**Figure S7, related to Figure 5**. **(A)** Western blot analysis of cell used for the RNA sequencing experiment validating depletion of ADNP one day or 14 days after treatment with 250nM dTAG13 (Control = DMSO) **(B)** Differential gene expression analysis between ADNP/ADNP2 perturbation and the corresponding unperturbed controls Significantly changing genes are colored (FDR < 0.01, |log2FoldChange| >1, n = 3) **(C)** Biological process GO term enrichment analyses of differentially changing gene sets for the indicated conditions. Note that there were no significantly enriched (FDR < 0.01) terms found for other category/condition combinations. **(D)** Distribution of distances from gene transcription start site to the nearest ADNP or ADNP2 peak split by regulation status in the combined ADNP/ADNP2 removal condition.

## MATERIALS AND METHODS

### Cell culture

Mouse embryonic stem cells (129 × C57BL/6 background) with BirA and Cre insertions in the Rosa26 locus^36,59^ were cultured on gelatin-coated dishes in ES medium containing DMEM (GIBCO 21969-035), supplemented with 15% fetal bovine serum (FBS; GIBCO), 1 × non-essential amino acids (GIBCO), 1 mM sodium pyruvate (GIBCO), 2 mM l-glutamine (GIBCO), 0.1 mM 2-mercaptoethanol (Sigma), 50 mg/ml penicillin, 80 mg/ml streptomycin, 3 μM glycogen synthase kinase (GSK) inhibitor (Calbiochem, D00163483 or Sigma, CHIR99021), 10 μM MEK inhibitor (Tocris, PD0325901), and homemade LIF, at 37°C in 5% CO2.

### Genome editing

Cells were trypsinized, counted, seeded, and immediately transfected using lipofectamine 3000 (Invitrogen) according to manufacturer instructions. Generally, 300k cells were seeded in 6-well plates and transfected with a total of 1-1.5mg DNA. For genetic knockout, plasmids encoding gRNAs targeting the N terminus and C terminus, Cas9 and a puromycin resistance cassette were co-transfected for 24h, before selection with 2mg/mL puromycin for a further 24-36h. For endogenous tagging, a plasmid harboring a desired homology repair cassette was included. Combined CRISPR-Cas9/TALEN editing was done as described previously^36^. The cells were then trypsinized, counted and 15000 cells were seeded on a 10cm dish for colony formation without puromycin selection. When colonies were sufficiently large (4-8 days), they were manually picked and split into two 96-well plates for screening and expansion. Screening for both positive editing events and unedited loci was performed by PCR and Sanger sequencing of the products. Where applicable, further confirmation was done using western blotting, RT-qPCR and RNA-sequencing.

### *In vitro* protein purification and analysis

For cloning, cDNA encoding full-length human ADNP2 (amino acid residues 1–1131) was PCR amplified and cloned into a pFast-Bac-derived vector (Invitrogen) in frame with an N-terminal His_6_-tag. An expression constructs encoding full-length human HP1β (amino acid residues 1– 185) was generated by amplification of cDNA and cloning into a pFast-Bac-derived vector in frame with an N-terminal Strep-tag II. cDNA encoding for full-length human CHD4 (amino acid residues 1–1912) was amplified and cloned into a pAC8-derived vector in frame with an N-terminal His_6_-tag^60^.

Baculoviruses for protein expression were generated in *Spodoptera frugiperda* Sf9 cells using the Bac-to-Bac method for pFastBac-derived vectors or by cotransfection with viral DNA for pAC8-based vectors. After one round of virus amplification in Sf9 cells, *Trichoplusia ni* High5 cells were infected with the respective Baculovirus (150 µl of virus per 10 ml of High5 cells at a density of 2 × 10^6^ cells ml^−1^) and collected 48 h after infection. Cells were lysed by sonication in 50 mM Tris, pH 7.5, 300 mM NaCl, 5 mM β-mercaptoethanol, 0.1% Triton X-100, 1 mM PMSF, 1× PIC (Sigma-Aldrich). The cleared lysate was passed over a Strep-Tactin Sepharose (IBA) column. The bound complex was eluted in 50 mM Tris-HCl, pH 7.5, 100 mM NaCl, 5 mM β-mercaptoethanol, 2.5 mM desthiobiotin.

### Analytical size-exclusion chromatography coupled to multi-angle light scattering (SEC-MALS)

Thirty eight microliters of sample at ~4-5 mg/mL of protein was injected onto a Superose 6 Increase 10/300 GL column (Cytiva) in 50 mM HEPES-OH, pH 7.4, 150 mM NaCl and 0.5 mM TCEP using an Agilent Infinity 1260 II HPLC system. In-line refractive index and light scattering measurements were performed using a Wyatt Optilab T-rEX refractive index detector and a Wyatt miniDAWN TREOS 3 light scattering detector. System control and analysis was carried out using the Wyatt Astra 7.3.1 software. System performance was checked with BSA.

### Western blotting

Protein samples in lauryl-dodecyl sulfate sample buffer (LDS) were separated by standard SDS-PAGE on 4-12% Bis-Tris gradient gels (Novex Bolt, Invitrogen). Separated proteins were transferred onto PVDF membranes (Milipore) in transfer buffer (Bjerrum-Schaeffer-Nielsen + 0.4% SDS) using a semi-dry transfer procedure (TransBlot Turbo, BioRad) with transfer parameters: 1.3A, 25V, 12min for one gel. The membranes were blocked in 3-5% skimmed milk (Sigma) dissolved in Tris buffered saline + 0.2% v/v Tween-20 (TBST). Antibody incubations were performed with antibodies diluted in blocking solution to empirically determined concentrations for a minimum of 1h up to a maximum of 24h. When re-probing, membranes were first treated with 0.01% NaN_3_ for 30-60min to quench the HRP from previous staining rounds. Between each antibody incubation, the membranes were washed for a minimum of 45min in TBST with at least 4 buffer exchanges. The horseradish peroxidase system (Immobilon, Milipore), coupled to camera-based detection (AI600, Agilent technologies) was used to visualize protein bands.

### Immune precipitations

Cells from one 10cm dish per condition were harvested by trypsinization before centrifugation (200gav, 5min). Cells were washed once in room temperature PBS and snap frozen or immediately taken forward for lysis. Pellets were lysed in NP-40 lysis buffer supplemented with protease inhibitors and benzonase (20mM Tris-HCl pH7.4, 150mM NaCl, 1% (v/v) NP-40, 0.1% (v/v) sodium deoxycholate,1xHALT protease inhibitor cocktail, 200U Turbo Benzonase (Milipore)) for 30min at 12°C and cleared by centrifugation at 4°C (16000gav, 20min). Protein concentration in the lysates was determined by Bradford assay against BSA and 1-3mg protein-lysate equivalent was loaded onto 10μL of bead slurry (Dynabeads, GE healthcare) prewashed twice with lysis buffer. For FLAG IPs, 2μg of antibody was added per 1mg of protein. Beads were incubated with lysate for 2h at 4°C, washed twice with lysis buffer and twice with wash buffer (20mM Tris-HCl pH7.4, 150mM NaCl, 0.1%(v/v) NP-40) before addition of LDS. All bead separation steps were done using magnetic racks.

### Proteomics

For IP-MS, immune precipitations were performed as normal with the addition of 2 washing steps without detergent (20mM Tris pH7.5, 150mM NaCl), followed by on-bead digestion. IP-MS in Figure 1B was exceptionally performed in 350mM NaCl, and Streptavidin M280 dynabeads (Invitrogen) used for pulldown, with all other steps remaining the same. Beads were resuspended by vortexing in 5 µL of digestion buffer (3M GuaHCl, 20mM EPPS pH8.5, 10mM CAA, 5mM TCEP) and 1 µL of 0.2 mg/mL LysC protease (Promega) in 50 mM HEPES (pH 8.5) was added. Proteins were digested for 2 h rotating at room temperature. The samples were diluted with 17 µL of 50 mM HEPES (pH 8.5) and digested with 1 µL of 0.2 mg/mL trypsin (Promega) in 0.2 mM HCl at 37°C with interval mixing at 2000 RPM for 30 sec every 15 min.

For Figure 1B, digested peptides were acidified with 0.8% TFA (final) and analyzed by LC– MS/MS on an EASY-nLC 1000 (Thermo Scientific) with a two-column setup. Peptides were applied on an Acclaim PepMap 100 C18 trap column (75 µm ID × 2 cm, 3 µm; Thermo Scientific) in 0.1% formic acid and 2% acetonitrile in H_2_O at a constant pressure of 80 MPa and separated by a linear gradient of 2%–6% buffer B in buffer A for 3 min, 6%–22% for 40 min, 22%–28% for 9 min, 28%–36% for 8 min, and 36%–80% for 1 min, and 80% buffer B in buffer A for 14 min (buffer A: 0.1% formic acid; buffer B: 0.1% formic acid in acetonitrile) on an EASY-Spray column ES801 (50 µm ID × 15 cm, 2 µm; Thermo Scientific) mounted on a DPV ion source (New Objective) connected to an Orbitrap Fusion (Thermo Scientific) at 150 uL/min flow rate. Data were acquired using 120,000 resolution for the peptide measurements in the Orbitrap and a top T (3-sec) method with HCD fragmentation for each precursor and fragment measurement in the ion trap following the manufacturer guidelines (Thermo Scientific).

For Figure S2, the following run parameters were used:

Instrument: Orbitrap Fusion LUMOS (Thermo Fisher Scientific) with VanquishNeo-nLC and an easy source with a 75um×15cm EasyC18 column. The samples were loaded on a C18 (0.3×5mm) trap and backward flush was used for the analysis. Gradient: 0-3min 2-4% buffer B in buffer A, 3-43min 4-20%, 43-58min 20-30%, 58-66min 30-36%, 66-68min 36-45%, 68-69min 45-100%, 69-75min 100% (buffer A: 0.1%FA in H2O; buffer B: 0.1%FA, 80% MeCN in H2O) at room temperature and the flow rate during the gradient was 350ul/min.

Peptides were identified with MaxQuant version 1.5.3.8 using the search engine Andromeda^61^. The mouse subset of the UniProt version 2017_04 or 2021_05 combined with the contaminant DB from MaxQuant was searched and the protein and peptide FDR values were set to 0.05. Statistical analysis was done in Perseus version 1.5.2.6^62^ or using limma within the einProt package (version 0.5.13)^63^. Results were filtered to remove reverse hits, contaminants and peptides found in only one sample. Missing values were imputed and potential interactors visualized in volcano plots.

### ChIP-seq

ChIP experiments were performed with at least 2 different clones from endogenously tagged cell lines. Harvesting was performed by trypsinization, and cells were counted for each sample. For ADNP2 ChIPs 2*10^7^ cells were collected, and for all other ChIPs 1*10^7^. The cells were crosslinked for 8 min at room temperature in 10ml PBS supplemented with 1% formaldehyde (Sigma, F8775). Cross-linking was quenched by adding glycine to a final concentration of 0.125 mM and incubating at room temperature for 1min, and on ice for 3min. Cells were pelleted by centrifugation at 500 g for 3 min at 4°C and the pellet was lysed in 10mL lysis buffer A (50 mM HEPES pH 8.0, 140 mM NaCl, 1 mM EDTA, 10% glycerol, 0.5% NP40, 0.25% Triton X-100) for 10 minutes on ice. After centrifugation the pellet was resuspended in 10 mL buffer B (10 mM Tris pH 8, 1 mM EDTA, 0.5 mM EGTA and 200 mM NaCl) and incubated for on ice for 5min. The samples were centrifuged at 500 g for 3 min at 4°C, and the pellets lysed in 180ul buffer C (50 mM Tris pH 8, 5 mM EDTA, 1% SDS, 100 mM NaCl) for 2min at room temperature and on ice for 10 min. The lysates were diluted in 1.6 mL ice cold TE buffer and sonicated in 15 mL tubes two times 10 cycles, 30 s ON / 30 s OFF, at 4°C (Bioruptor Pico, BioRad). Then, 200ul 10x ChIP buffer (0.1% SDS, 10% Triton X-100, 12mM EDTA, 167mM Tris-HCl pH 8, 1.67M NaCl) was added, and the chromatin was transferred into 2 mL Eppendorf tubes before centrifugation for 10 min at 16000 g, 4°C. 5% sheared chromatin was reserved for the input control, while the rest was transferred into fresh tubes. Generally, beads were prewashed in 1x ChIP buffer, and added to the sonicated chromatin, alongside different amounts of antibody, depending on the ChIP. For ADNP2, 20uL Protein G Dynabeads and 2uL a-FLAG. For H3K9me3, 20uL Protein G Dynabeads and 2uL a-H3K9me3. For ADNP, 20uL Protein G Dynabeads and 20uL Protein A Dynabeads were pre-mixed and washed twice with 1x ChIP buffer, and 3uL of a-FLAG used. Samples were incubated for 4h at 4°C. ChIPs were washed for 1 minute each for each step, 4 times RIPA (10mM Tris-HCl pH 8.0, 1mM EDTA pH 8.0, 140mM NaCl, 1% Triton X-100, 0.1% SDS, 0.1% Na-deoxycholate), 2 times RIPA500 (10mM Tris-HCl pH 8.0, 1mM EDTA pH 8.0, 500mM NaCl, 1% Triton X-100, 0.1% SDS, 0.1% Na-deoxycholate), 2 times Li-wash buffer (10mM Tris-HCl pH 8.0, 1mM EDTA, pH 8.0, 250mM LiCl, 0.5% NP-40, 0.5% Na-deoxycholate), 1x TEplus (10mM Tris-HCl pH 8.0, 1mM EDTA). Beads were transferred to a fresh tube during the last wash and wash buffer was completely removed before adding 75uL elution buffer ((10mM Tris-HCl pH 8.0, 1mM EDTA pH 8.0, 150mM NaCl, 1% SDS) and incubating 20 minutes at 65°C with constant shaking. Elution was repeated once more with 75uL elution buffer for 20 minutes and eluates were pooled, 2 ul RNaseA (20μg/ul) were added and incubated for 1 h at 37°C. Then 2ul Proteinase K (20mg/ml) was added and samples were incubated 2 h at 55°C followed by decrosslinking for 6 h at 65°C. Input samples were adjusted to 150ul total volume with elution buffer and processed equivalently to ChIP samples. DNA was purified by adding 30ul AMPure XP beads, 9 ul 5M NaCl, and 190ul Isopropanol and incubating for 10 min at RT after thorough mixing. The beads were collected on a magnetic rack, washed twice with 80% EtOH and DNA was eluted in 30ul 10mM Tris pH8.0 for 5 min at 37°C. 25ul ChIP DNA or 10ng Input DNA were used to generate libraries using the NEBNext Ultra II Library Prep Kit for Illumina (NEB). Reactions were scaled down to half otherwise processing was according to the manufacturer’s manual. Libraries were sequenced 51bp paired-end on a NovaSeq6000 instrument (Illumina), 75bp paired-end on a NextSeq2000 device (Ilumina), or 50bp single-end on HiSeq2500 (Ilumina).

### ATAC-seq

ATAC-seq was performed in biological triplicates as previously described^36^.

### RNA-seq

RNA was prepared equally as for RT-qPCR. Libraries were prepared using the Ilumina Stranded Total RNA Library Prep, including a ribosomal RNA depletion step, and sequenced on the Illumina NovaSeq 6000 (51-nt paired-end reads).

### RT-qPCR

RNA was isolated from cells using the Absolutely RNA miniprep kit (Agilent) according to the manufacturer’s instructions, including genomic DNA pre-filtering and DNaseI treatment. The concentration was determined with the RNA Broad Range reagents on a Qubit 2.0 system according to the manufacturer’s instruction. Reverse transcription was performed by adding Primescript II master mix (Takara) to 1x concentration and incubating for 15min at 37°C, followed by enzyme inactivation at 85°C for 5s. qPCR was performed with the SSO advanced BioRad qPCR master mix using an amount of inactivated RT mixture corresponding to 40ng-200ng total RNA (depending on experiment), with 0.4uM primers, in a CFX96 system (BioRad). The cycling parameters were always: 30s 95°C, 40 cycles [5s 95°C, 15s 60°C], melt curve 65°C-95°C.

### Computational methods

#### Read mapping

Reads were mapped to the mouse mm10 genome or a combined mouse/human (mm10/GRCh38) genome (for the samples including spike-ins) using STAR version 2.7.3^64^ (allowing up to 10000 multi-mapping reads, reporting 1 multi-mapper at a random location).

ChIP: STAR --runMode alignReads --outSAMtype BAM SortedByCoordinate --readFilesIn R1.fastq.gz R2.fastq.gz --readFilesCommand zcat --genomeDir mm10_hg38Spike_refSTAR --runThreadN 10 --alignIntronMax 1 --alignEndsType EndToEnd --outFilterType Normal --seedSearchStartLmax 30 --outFilterMultimapNmax 10000 --outSAMattributes NH HI NM MD AS nM --outMultimapperOrder Random --outSAMmultNmax 1 --outSAMunmapped Within --outFileNamePrefix _ --clip3pAdapterSeq CTGTCTCTTATACACATCT AGATGTGTATAAGAGACAG --outBAMsortingBinsN 100

RNAseq: STAR --runMode alignReads --outSAMtype BAM SortedByCoordinate --readFilesIn R1.fastq.gz R2.fastq.gz --readFilesCommand zcat –genomeDir tar2_7_3a_GRCm38.primary_assembly_gencodeM23_spliced_sjdb50 --runThreadN 10 --outFilterType BySJout --outFilterMultimapNmax 10000 --outFilterMismatchNmax 3 --winAnchorMultimapNmax 20000 --alignMatesGapMax 350 --seedSearchStartLmax 30 alignTranscriptsPerReadNmax 30000 --alignWindowsPerReadNmax 30000 --alignTranscriptsPerWindowNmax 300 --seedPerReadNmax 3000 --seedPerWindowNmax 300 --seedNoneLociPerWindow 1000 --outSAMattributes NH HI NM MD AS nM --outMultimapperOrder Random --outSAMmultNmax 10000 --outSAMunmapped Within

We used RepeatMasker (options: -species “Mus musculus”) (A.F.A. Smit, R. Hubley & P. Green RepeatMasker at http://repeatmasker.org) to annotate repeats and extract consensus repeat sequences. Reads that overlap repeat annotations were extracted and aligned to repeat consensus sequences using bowtie2^65^ (version: 2.3.5.1, options: -q -D 20 -R 3 -N 1 -L 20 -i S,1,0.50 --local -p 20 --no-unal).

#### Identification and annotation of ADNP2/ADNP binding sites (Peak finding)

To identify peaks in ADNP2 or ADNP ChIPs, we pooled all ChIP replicates and used the callpeak function of MACS2^66^ (version 2.2.7.1). Peaks were resized to span 300bp around the peak summit. To filter out only significantly enriched peaks, we kept only those peaks where the ChIP was enriched > 1.2 fold over input in at least 3 replicates.

We used Gencode version M23 annotation and defined transcription start sites (TSSs) as 300bp upstream of the annotated TSS until the TSS. To find overlaps between peaks and TSSs or repeat elements, we used the Genomic Ranges R package^67^. We also generated 100 sets of randomly distributed regions matching our peak sets in number and size, and calculated overlaps with TSS and repeat annotations using these regions. Repeat annotation overlaps were summed up based on the repeat_name column of the repeat masker output (generated as described above).

#### ChIP-seq analysis

To determine differences in ADNP2 or ADNP ChIP signals in WT and mutant cells, we used Quasr^68^ to count the number of reads in peaks or repeat elements and edgeR^69^ for differential count analysis and normalization to library size (using, if available, the human spike-in total read counts as library size, TMM normalization, and a prior.count of 3). To display ChIP signal intensities (as counts per million) in heatmaps or metaplots, or to calculate cpm in sliding windows across a chromosome, we used the MiniChip R package (https://github.com/fmi-basel/gbuehler-MiniChip). We used MEME-ChIP version 5.5.5^70^ to search for motifs in all 6315 ADNP2 peaks (using the sequence 500bp around the peak summit and the option -maxw 25).

#### RNA-seq analysis

To count the number of uniquely mapping reads in genes, we used featureCounts from the Rsubread^71^ package on GENCODE version M24 gene annotation. We used TEtranscripts^72^ to quantify read counts within transposable elements (TEs) with a TE annotation for mm10 downloaded from https://labshare.cshl.edu/shares/mhammelllab/www-data/TEtranscripts/TE_GTF/mm10_rmsk_TE.gtf.gz and the GENCODE version M24 gene annotation and the options --format BAM --sortByPos --stranded reverse --mode multi.

We then calculated differential expression and counts per million using DESeq2^73^.

To find enriched gene ontology terms among regulated genes we used the enrichGO function of the clusterProfiler^74^ R package.

## Data and Code Availability

All NGS data generated for this paper has been deposited at NCBI GEO and are available under accession number GEO: GSE253069. Custom scripts for data analysis are available on GitHub (https://github.com/xxxmichixxx/ChAHP2). The mass spectrometry proteomics data have been deposited to the ProteomeXchange Consortium via the PRIDE^75^ partner repository with the dataset identifiers PXD048314 and PXD048310. Other tools used are indicated in the respective Method Details sections.

